# Integrated Resistome and Quantitative Proteomics Reveal Coordinated Resistance Architecture in MDR and XDR Gram-Negative ICU Pathogens

**DOI:** 10.64898/2026.04.15.718841

**Authors:** Aldo A.M. Lima, Dilza T. Silva, Nicholas E. Sherman, Lavouisier F.B. Nogueira, Marília S. Maia, Marco A.F Clementino, Alexandre Havt, Jose Q.S. Filho, S. Francisco, Ila F.N. Lima, Deiziane V.S. Costa, Samilly A. Ribeiro, Felipe P. Mesquita, José K. Sousa, Luis F. B. Lino, Ana E. S. Alves, Andreza C. Damasceno, Lyvia M.V.C. Magalhães, Rafhaella N.D.G. Gondim, Luciana V.C. Fragoso, Jorge L.N. Rodrigues, Fabio Miyajima, Bráulio M. Carvalho, Érico A.G. Arruda

## Abstract

**Objectives:** Antimicrobial resistance (AMR) in Gram-negative pathogens is driven by complex and coordinated molecular mechanisms that remain incompletely characterized. This study integrated phenotypic, genomic, and quantitative proteomic analyses to characterize multidrug-resistant (MDR) and extensively drug-resistant (XDR) Gram-negative bacteria circulating in an intensive care unit (ICU) in Northeastern Brazil.

**Methods:** A total of 259 Gram-negative isolates collected between 2019 and 2021 underwent species identification, antimicrobial susceptibility testing, and targeted qPCR for resistance genes. *Klebsiella pneumoniae*, *Acinetobacter baumannii*, and *Pseudomonas aeruginosa* representing susceptible, MDR, and XDR phenotypes were selected for whole-genome sequencing and label-free quantitative proteomics. Differential protein abundance was assessed using Limma with |log2FC| > 1 and p < 0.05.

**Results:** *K. pneumoniae* (47%), *A. baumannii* (24%), and *P. aeruginosa* (21%) predominated. Carbapenem resistance reached 44%, 93%, and 61%, respectively, and MDR/XDR phenotypes occurred in >30% of isolates. Genomic analyses revealed dense resistomes with coexisting β-lactamases (blaKPC, blaNDM, blaCTX-M, OXA) and widespread efflux systems. Proteomic profiling demonstrated phenotype-associated differences in outer membrane proteins, transport systems, regulatory proteins, and metabolic pathways. XDR isolates showed additional enrichment of envelope remodeling proteins, stress response mechanisms, and proteostasis-associated factors.

**Conclusions:** MDR and XDR Gram-negative ICU pathogens exhibit coordinated resistance architecture characterized by accumulation of resistance genes and adaptive proteomic remodeling. Integrated multi-omics approaches provide mechanistic insight into antimicrobial resistance and support improved surveillance and therapeutic strategies.

**What is known?:**

- Antimicrobial resistance is a priority and a serious problem in global health, resulting in high rates of morbidity and mortality.
- *Klebsiella pneumoniae*, *Acinetobacter baumannii*, and *Pseudomonas aeruginosa* are on the World Health Organization’s (WHO) priority list as major causes of morbidity and mortality worldwide.
- Classical characterization of susceptibility and resistance phenotypes does not capture the complexity of antimicrobial resistance and hampers effective control measures and actions to minimize the evolutionary dynamics of resistance in these bacteria.

**What is new?:**

- The study characterizes the phenotypic pattern of antimicrobial susceptibility, the presence and sequencing of the resistome and virulome, and analyzes the label-free quantitative proteome of susceptible, MDR, and XDR phenotypes in strains of *K. pneumoniae*, *A. baumannii*, and *P. aeruginosa* circulating in hospital ICUs in Brazil.
- MDR and XDR gram-negative phenotypes are associated with a dense resistome, with widespread dissemination of beta-lactamase genes (*bla_KPC*, *bla_NDM*, *bla_CTX-M*, and *OXA*) and RND-type (*MEX*s) and *acrAB-tolC* efflux pumps, without changes in virulence genes.
- Proteomic analysis demonstrated increased production of beta-lactamases, components of efflux pump systems, outer membrane protein synthesis, protection for oxidative stress mechanisms, proteins for iron acquisition, and systemic regulators. XDR strains additionally showed enhanced remodeling of the cell envelope, activation of proteostasis, and metabolic adaptation.

## Introduction

Antimicrobial resistance (AMR) is a major determinant of adverse clinical outcomes in bacterial infections. A recent global systematic analysis estimated that bacterial AMR was directly responsible for approximately 1.14 million deaths in 2021, with 4.95 million deaths associated with drug-resistant infections, reflecting both direct and indirect effects of resistance on patient outcomes (1,2). These findings underscore the substantial contribution of AMR to global mortality. Modelling forecasts suggest that, if current trends continue without effective intervention, drug-resistant infections could result in 39 million deaths worldwide between 2025 and 2050, with nearly 10% projected in Latin America and the Caribbean, emphasizing the urgency of the problem in Brazil and neighboring regions (2).

Beyond increased mortality, AMR is associated with prolonged infections, longer hospital stays, higher rates of intensive care unit (ICU) admission, and restricted therapeutic options (1). WHO GLASS surveillance data indicates persistently high resistance rates among Gram-negative pathogens, particularly in bloodstream and urinary tract infections (1). Gram-negative bacteria resistant to critical antimicrobials, including carbapenem-resistant *Klebsiella pneumoniae*, *Escherichia coli*, and *Acinetobacter baumannii*, represent a major clinical concern due to elevated case fatality rates and limited treatment options (3–5). Carbapenem-resistant *A. baumannii* (CRAB) has been associated with mortality rates ranging from 30% to 75% in critically ill patients (4,5). Among these pathogens, *Pseudomonas aeruginosa* exemplifies adaptability and persistence due to its large genome, regulatory complexity, and capacity to thrive under environmental stress (6,7,8). The burden of AMR is disproportionately higher in settings with limited diagnostic and surveillance capacity, where delayed or inappropriate therapy further increases morbidity and mortality (1,4).

Brazil accounts for a substantial AMR burden in Latin America, with approximately 33,200 deaths directly attributable to bacterial AMR annually and 137,900 deaths associated with resistant infections each year (6). Multidrug-resistant Gram-negative bacteria have emerged as leading causes of severe healthcare-associated infections, particularly in ICUs, where antimicrobial pressure, invasive procedures, and vulnerable hosts converge (3,7).

Resistance mechanisms include low outer membrane permeability, porin modulation, overexpression of multidrug efflux pumps (e.g., MexAB-OprM), enzymatic inactivation via β-lactamases, and rapid adaptation through mutations and horizontal gene transfer (8–10). Emerging resistance to novel β-lactam/β-lactamase inhibitor combinations has been linked to structural mutations in AmpC, OXA-type enzymes, carbapenemases, efflux regulators, and iron-uptake systems (9,10). Biofilm formation and adaptive proteomic responses further enhance antimicrobial tolerance and persistence in critically ill patients (8).

We hypothesized that MDR, XDR, and carbapenem-resistant Gram-negative bacteria circulating in ICUs in Northeastern Brazil harbor coordinated resistance architectures driven by accumulation of β-lactamase genes, efflux pump operons, porin alterations, and adaptive proteomic remodeling (9,11–13).

## Material and Methods

### Study Design, Setting, and Ethical Approval

This study was a prospective held from January 2019 and December 2021 Gram-negative bacteria were isolated from patients admitted to the intensive care unit (ICU) of the Hospital Universitário Walter Cantídeo, a tertiary public hospital in Fortaleza, Ceará, Northeastern Brazil. The study was approved by the Brazilian National Research Ethics Committee (CAAE n° 03300218.2.0000.5054; approval n° 3.143.258). Informed consent was waived because analyses were performed on de-identified bacterial isolates collected as part of routine care.

Gram-negative isolates recovered during routine microbiological diagnostics were included in a laboratory-based cohort study integrating phenotypic antimicrobial susceptibility testing with genomic and quantitative proteomic analyses. Isolates were obtained from patients with suspected infection or colonization and originated from tracheal aspirates, bronchoalveolar lavage, blood cultures, urine, surgical wounds, sterile body fluids, tissue fragments, and rectal swabs collected for surveillance. Only one isolation per patient per species was included.

### Bacterial Identification and Antimicrobial Susceptibility Testing

Clinical specimens were processed according to standard microbiological procedures. Blood cultures were incubated in the BacT/ALERT® 3D system (bioMérieux). Non-sterile specimens were cultured on appropriate selective and non-selective media and incubated at 37 ± 2°C for 18–24 hours. Species identification and antimicrobial susceptibility testing were performed using the VITEK® 2 Compact automated system (bioMérieux). Minimum inhibitory concentrations (MICs) were interpreted according to CLSI criteria (April 2019–January 2020) and BrCAST criteria thereafter. Resistance phenotypes were classified as susceptible (SUSC), multidrug-resistant (MDR), extensively drug-resistant (XDR), or carbapenem-resistant (CR) according to international consensus definitions (14). Isolates of *Klebsiella pneumoniae*, *Acinetobacter baumannii*, and *Pseudomonas aeruginosa* were prioritized for molecular and proteomic analyses due to their clinical relevance and high resistance burden.

### DNA Extraction and Targeted Molecular Screening

Selected isolates were recovered from frozen stocks and cultured overnight in tryptic soy broth at 37°C with agitation. Genomic DNA was extracted using the Wizard® Genomic DNA Purification Kit (Promega). DNA concentration and purity were assessed by NanoDrop™ spectrophotometry and stored at −80°C. Targeted real-time PCR (qPCR) was performed using a QuantStudio™ 3 system (Applied Biosystems).

Primers were designed to cover major β-lactamase families, including: (a) Ambler class A: *bla_KPC (19 variants)*, *bla_CTX-M (142 variants)*, *bla_TEM (167 variants)*, *bla_SHV*, *bla_GES (25 variants)*; (b) Ambler class B: *bla_NDM (27 variants)*; and (c) Ambler class D: OXA-type genes (203 variants including *OXA-23-like*, *OXA-24/40-like*, *OXA-48-like*, *OXA-51-like*). Primer design incorporated conserved regions to maximize variant coverage while preserving specificity. In parallel, primers targeting 12 genes within Resistance–Nodulation–Division (RND) efflux systems (*mex* operons) were developed.

Reactions included positive controls (isolated from confirmed resistant isolations) and negative controls. Cycling conditions consisted of initial denaturation (95°C, 2 min) followed by 35 cycles of denaturation (95°C, 15 s) and annealing/extension (primer-specific temperature, 1 min). Melting curve analysis confirmed amplification specificity. Molecular findings were integrated with phenotypic resistance profiles.

### Whole-Genome Sequencing and Bioinformatic Analysis

Representative isolates spanning SUSC, MDR, and XDR phenotypes underwent whole-genome sequencing (WGS). Genomic libraries were prepared using the Illumina DNA Prep kit in a miniaturized format. Library preparation was automated using Dragonfly Discovery and Mosquito liquid handling systems (SPT Labtech), enabling 384-sample batch processing. DNA underwent tagmentation, magnetic bead purification, indexing, amplification, and size selection using SPRI Select beads (Beckman Coulter). Libraries were pooled (4 × 96 samples), quantified using Qubit™ fluorimetry, and sequenced on the Illumina NextSeq 1000/2000 platform with paired end reads. FASTQ files were processed using the Bactopia pipeline on the IAM Carlos Chagas computational cluster (Fiocruz-PE). De Novo assembly was performed with SPAdes. Assembly quality was evaluated using N50, contig number, and coverage metrics. Resistance genes, virulence determinants, and plasmid replicons were identified using ABRicate with the following databases: (a) ResFinder (antimicrobial resistance genes); (b) VFDB (virulence genes); and (c) PlasmidFinder (plasmid typing). Automated annotations were manually curated to remove redundancies and confirm gene classification.

### Quantitative Proteomic Analysis

Nine representative strains (three SUSC, three MDR, three XDR) across the three species were selected for label-free quantitative proteomics. Proteomic analyses were performed at the Biomolecular Analysis Facility (BCA, University of Virginia). Cell pellets were disrupted using bath sonication, tip sonication, and bead beating. Protein concentration was determined by BCA assay. 30 micrograms of protein per sample were processed using the PREOMICS iST 96X HT kit following manufacturer protocols. Peptide cleanup was performed using ZipTip C18 columns. Peptide quantification was performed fluorometrically, and 500 ng were injected per analysis.

Samples were analyzed on an Orbitrap Exploris 480 (Thermo Fisher Scientific). Raw data were searched using Proteome Discoverer (v2.5.0.400) against UniProt species-specific databases. Trypsin/P was specified as the cleavage enzyme (one missed cleavage allowed). Mass tolerances were set at 10 ppm (precursor) and 0.02 Da (fragment ions). Carbamidomethylation of cysteine was fixed; methionine oxidation was variable.

Protein identification and quantification were validated using Scaffold Q+S (v5.3.4).

### Statistical and Bioinformatic Analysis

#### Phenotypic and Genomic Data

Categorical variables were expressed as frequencies; continuous variables as medians and interquartile ranges (IQR). Comparisons across phenotypic groups were performed using Fisher’s exact test or Kruskal–Wallis’s test, as appropriate. Odds ratios (ORs) with 95% confidence intervals (CI) were calculated for associations between resistance genes and phenotypes. A two-sided p-value < 0.05 was considered statistically significant. Analyses were conducted using SPSS v21.0.

#### Proteomic Data

Label-free quantification was performed using total ion current normalization. Protein intensities were log₂-transformed. Proteins with missing values across samples were filtered out. Differential protein abundance was assessed using the Limma package in R, applying linear models with empirical Bayes moderation. Statistical thresholds for differential enrichment were defined within the Limma framework. Data visualization, including volcano plots and boxplots, was generated using ggplot2. Functional interpretation focuses on antimicrobial inactivation, efflux systems, envelope remodeling, oxidative stress responses, iron acquisition pathways, and global regulatory proteins.

#### Study Integration Framework

All analyses were exploratory and designed to integrate phenotypic resistance patterns with molecular and proteomic data. The objective was to characterize associations between resistance phenotypes, resistome complexity, virulence gene content, and protein expression signatures in Gram-negative ICU isolates.

## Results

### Study Flow and Isolate Selection

A total of 259 Gram-negative bacterial isolates were recovered from patients admitted to the intensive care unit (ICU) of a tertiary public hospital in Fortaleza, Northeastern Brazil, between 2019 and 2021. Isolates were obtained from routine clinical and surveillance specimens, and only one isolate per patient per species was included in the study. All isolates underwent species identification and antimicrobial susceptibility testing. Phenotypes were classified as susceptible (SUSC), multidrug-resistant (MDR), extensively drug-resistant (XDR), or carbapenem-resistant (CR) according to international consensus definitions (14). Out of the 259 isolates, *Klebsiella pneumoniae* (n = 122), *Acinetobacter baumannii* (n = 61), and *Pseudomonas aeruginosa* (n = 54) were prioritized for molecular and proteomic analyses due to their clinical relevance and high resistance burden. The overall workflow integrating phenotypic, molecular, genomic, and proteomic analyses is summarized in **Figure 1**.

**Figure 1.**
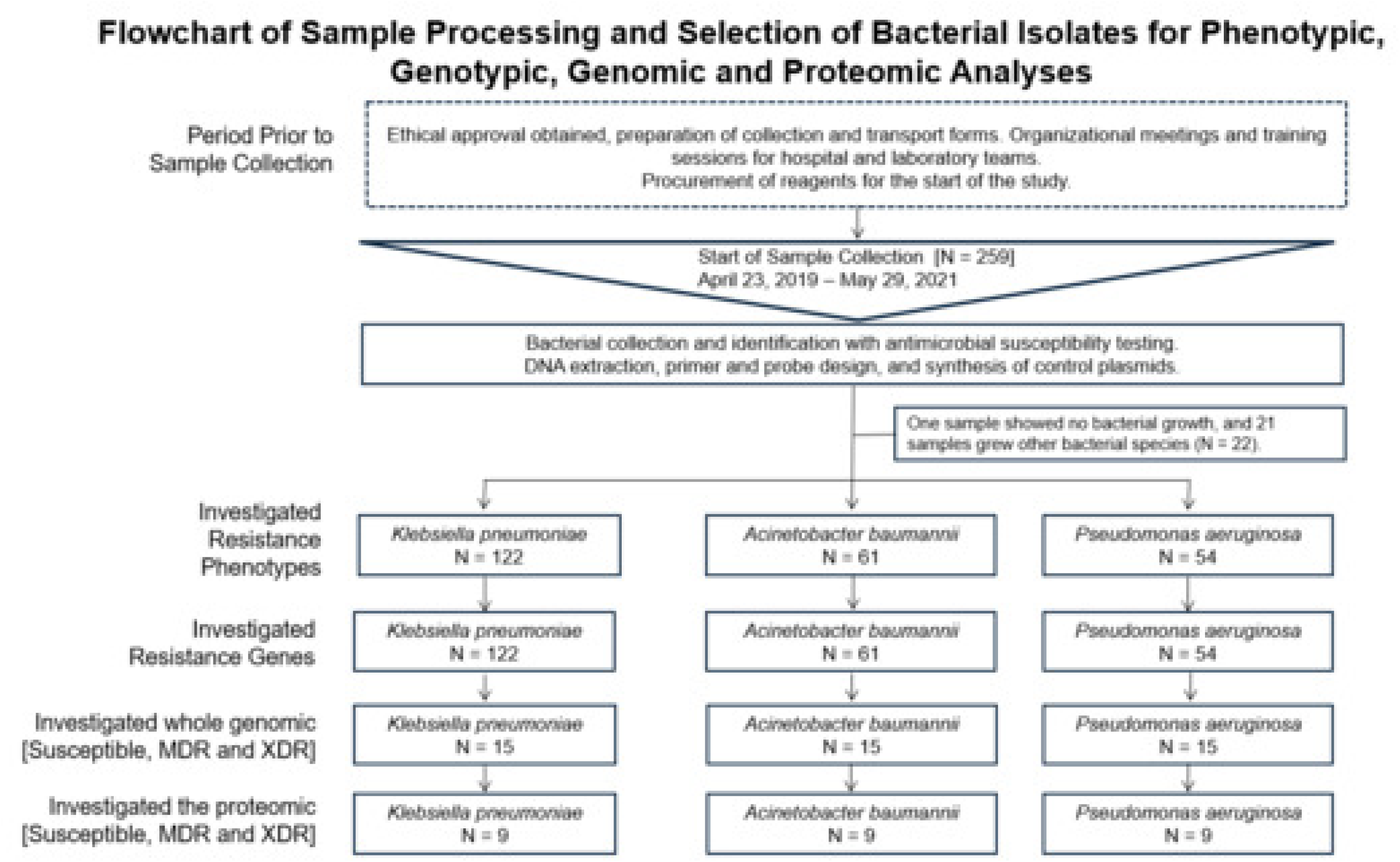
Flowchart of the study design and isolate selection. Flow diagram illustrating the selection of Gram-negative bacterial isolates recovered from patients hospitalized in the intensive care unit (ICU) in Fortaleza, Ceará, Brazil, between 2019 and 2021. The figure summarizes the total number of clinical specimens screened, isolates identified at the species level, exclusion criteria (e.g., duplicates or incomplete records), and the final number of isolates included for phenotypic antimicrobial susceptibility testing, whole-genome sequencing, and proteomic analysis.

### Clinical and Microbiological Characteristics

Among the 259 isolates, *K. pneumoniae* accounted for 47% (122/259), followed by *A. baumannii* (24%; 61/259) and *P. aeruginosa* (21%; 54/259) (**Figure 2**). The remaining 8% comprised other Enterobacterales and non-fermenters. Most isolates were recovered from male patients (65%), with a median age of 57 years (Internquartile range; IQR 48–70) (**Table 1**). Tracheal aspirates were the most common source of isolation (45%), followed by rectal swabs (26%) and blood cultures (12%). Species-specific distributions differed: *A. baumannii* and *P. aeruginosa* predominated in respiratory specimens (61% and 74%, respectively), whereas *K. pneumoniae* was frequently isolated from rectal swabs (53%) (**Figure 2** and **Table 1**). Hospital-acquired infections represented a substantial proportion of isolates, particularly among *A. baumannii* (41%) and *P. aeruginosa* (20%), consistent with their known association with ventilator-associated pneumonia and device-related infections (1,2).

**Figure 2.**
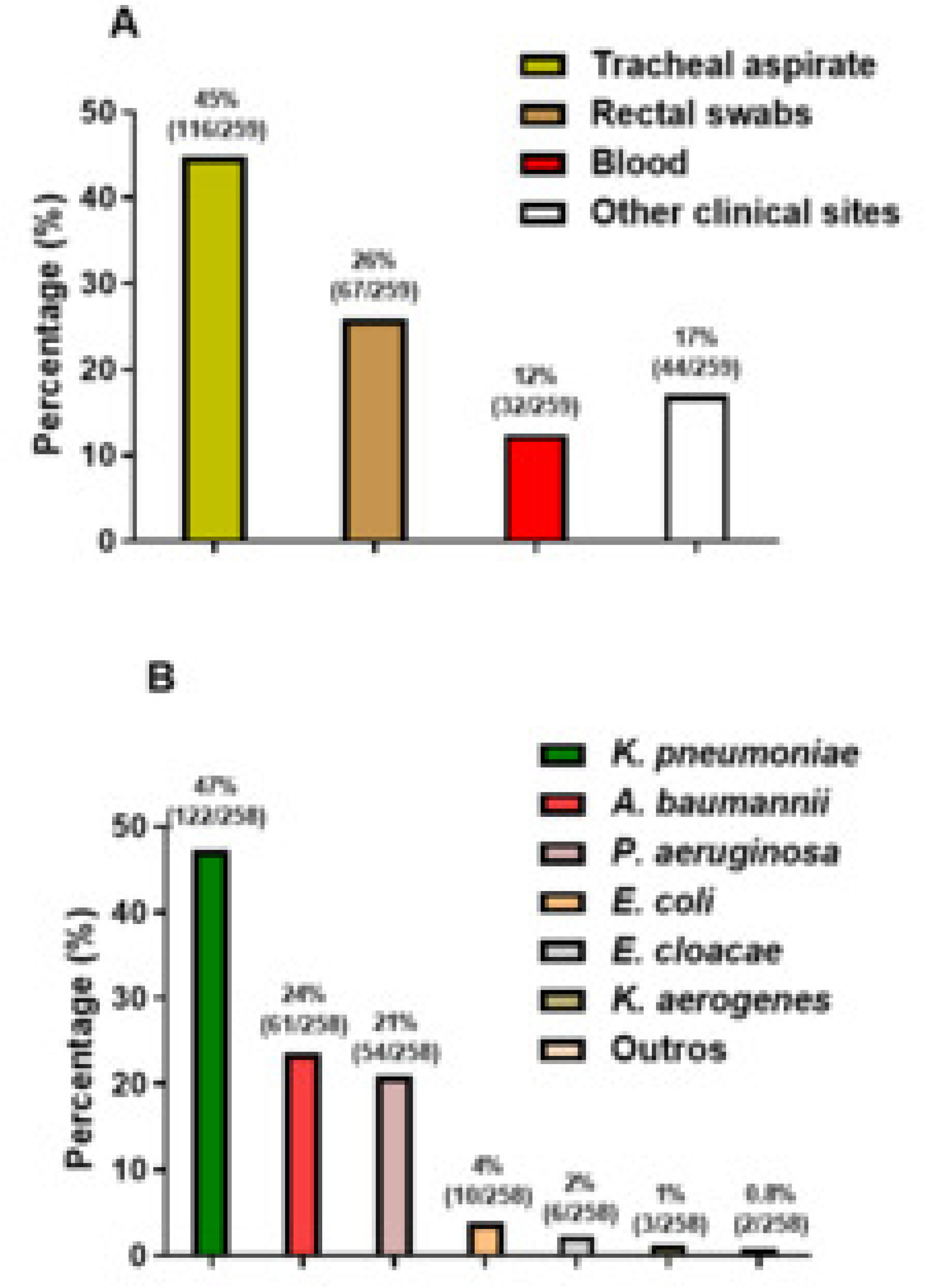
Distribution of clinical specimens and Gram-negative bacterial species isolated from ICU patients. (**A**) Proportion of clinical specimens collected, including tracheal aspirates, rectal swabs, blood, and other clinical sites. Percentages and absolute numbers (n/N) are indicated above each bar. (**B**) Distribution of Gram-negative bacterial species identified among the isolates, including *Klebsiella pneumoniae*, *Acinetobacter baumannii*, *Pseudomonas aeruginosa*, *Escherichia coli*, *Enterobacter cloacae*, *Klebsiella aerogenes*, and other species. Percentages and absolute numbers (n/N) are shown above each bar.

**Table 1-.**
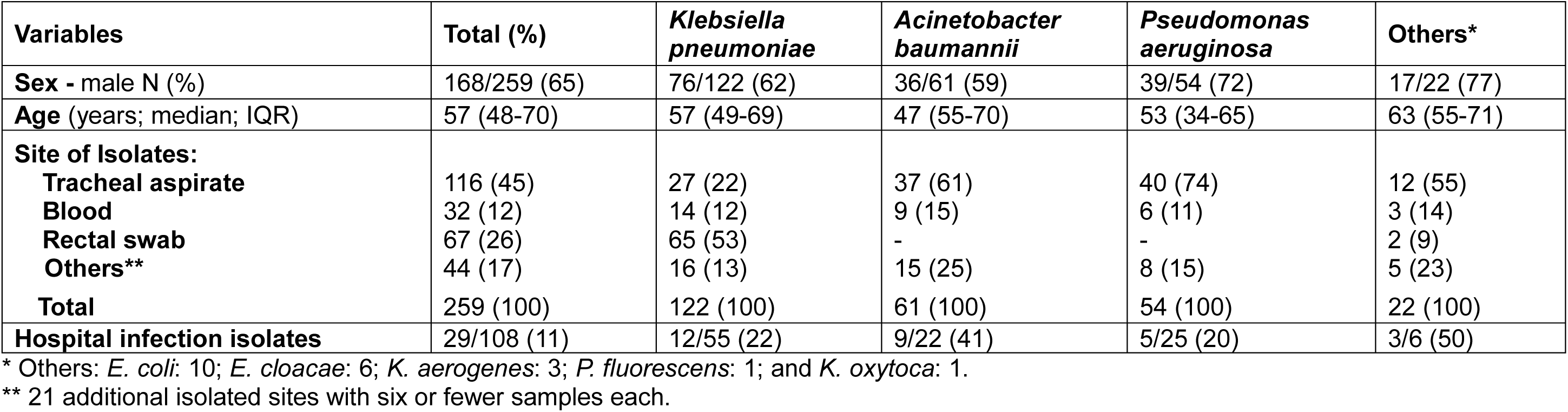
Characterization of Gram-Negative Bacterial Isolates from Patients Hospitalized in the Intensive Care Unit in Fortaleza, Ceará, Brazil, from 2019 to 2021.

| Variables | Total (%) | *Klebsiella pneumoniae* | *Acinetobacter baumannii* | *Pseudomonas aeruginosa* | Others* |
| --- | --- | --- | --- | --- | --- |
| **Sex** - male N (%) | 168/259 (65) | 76/122 (62) | 36/61 (59) | 39/54 (72) | 17/22 (77) |
| **Age** (years; median; IQR) | 57 (48-70) | 57 (49-69) | 47 (55-70) | 53 (34-65) | 63 (55-71) |
| **Site of Isolates:** |  |  |  |  |  |
| Tracheal aspirate | 116 (45) | 27 (22) | 37 (61) | 40 (74) | 12 (55) |
| Blood | 32 (12) | 14 (12) | 9 (15) | 6 (11) | 3 (14) |
| Rectal swab | 67 (26) | 65 (53) | - | - | 2 (9) |
| Others** | 44 (17) | 16 (13) | 15 (25) | 8 (15) | 5 (23) |
| **Total** | 259 (100) | 122 (100) | 61 (100) | 54 (100) | 22 (100) |
| **Hospital infection isolates** | 29/108 (11) | 12/55 (22) | 9/22 (41) | 5/25 (20) | 3/6 (50) |

\* Others: *E. coli*: 10; *E. cloacae*: 6; *K. aerogenes*: 3; *P. fluorescens*: 1; and *K. oxytoca*: 1.
\*\* 21 additional isolated sites with six or fewer samples each.

### Antimicrobial Resistance Phenotypes

A high burden of antimicrobial resistance was observed across all species, particularly against β-lactams, carbapenems, aminoglycosides, and fluoroquinolones (**Table Suppl. 1**). Our results show that resistance to β-lactam antibiotics was widespread. In *K. pneumoniae*, resistance exceeded 80% for most penicillins and cephalosporins, including cefepime (83%), ceftazidime (87%), ceftriaxone (83%), and cefuroxime (90%). Carbapenem resistance was also high, reaching 70% for meropenem and 72% for imipenem. *A. baumannii* exhibited the most extreme resistance profile. Resistance to penicillins and cephalosporins approached 100%, and carbapenem resistance was nearly universal (98% for meropenem; 93% for imipenem). *P. aeruginosa* demonstrated high resistance to extended-spectrum cephalosporins and carbapenems, with resistance rates of 56% to meropenem and 61% to imipenem. Extended-spectrum β-lactamase (ESBL) phenotypes were frequent, particularly in *A. baumannii* (93%) and *P. aeruginosa* (59%), and in nearly half of *K. pneumoniae* isolates (47%). Resistance to aminoglycosides varies across species. In *K. pneumoniae*, resistance to gentamicin was 57%, whereas amikacin resistance was lower (14%). *A. baumannii* exhibited resistance rates of 33% to gentamicin and 23% to amikacin. In *P. aeruginosa*, resistance reached 42% for gentamicin and 28% for amikacin.

Although aminoglycosides retained partial activity against selected isolates, effectiveness was markedly reduced in MDR and XDR phenotypes. Fluoroquinolone resistance was highly prevalent. Resistance to ciprofloxacin was detected in 91% of *K. pneumoniae* and 96% of *A. baumannii* isolates. In *P. aeruginosa*, 55% of isolates were resistant. Integration across antimicrobial classes demonstrated a predominance of MDR and XDR phenotypes. MDR phenotypes were identified in 37% of *K. pneumoniae*, 95% of *A. baumannii*, and 61% of *P. aeruginosa* isolates. XDR phenotypes affected 32%, 80%, and 54% of isolates, respectively. Carbapenem resistance was especially prominent in *A. baumannii* (93%) and *K. pneumoniae* (72%). Collectively, these findings indicate an ICU ecosystem dominated by high-level multidrug resistance with limited therapeutic options.

### Distribution of β-Lactamase and Efflux Pump Genes

Targeted molecular screening revealed widespread and heterogeneous distribution of β-lactamase and efflux pump genes (**Table Suppl. 2**). In *K. pneumoniae*, *bla_TEM* and *bla_SHV* were detected in 81% and 95% of isolates, respectively, and *bla_GES* in 21%. ESBL genes of the *bla_CTX-M* family were highly prevalent (>80% for major variants). Similar patterns were observed in *A. baumannii* and *P. aeruginosa*, indicating broad horizontal dissemination of ESBL determinants. Carbapenemase genes were strikingly common. In *K. pneumoniae*, *bla_KPC* was detected in 97% of isolates and *bla_NDM* in 34%, with co-occurrence in 34%. In *A. baumannii*, *bla_KPC* and *bla_NDM* were present in 86% and 36%, respectively. In *P. aeruginosa*, both genes were detected in 82% of isolates. *A. baumannii* showed high prevalence of *bla_OXA-23* (78%) and intrinsic *bla_OXA-51* (95%). OXA variants were less frequent in *K. pneumoniae* and *P. aeruginosa*. Genes encoding Resistance–Nodulation–Division (RND) efflux pumps were nearly ubiquitous. In *K. pneumoniae*, mex-associated genes were detected in high frequencies (e.g., *mexA* 100%, *mexB* 91%), and *acrAB-tolC* was present in 95%. In *A. baumannii*, mex genes were also frequent, though *acrAB-tolC* was rare (2%), reflecting species-specific efflux architectures. All *P. aeruginosa* isolates harbored complete and redundant mex operon repertoires (approaching 100% detection across multiple operons), consistent with its intrinsic multidrug resistance capacity. The coexistence of multiple β-lactamase classes with dense efflux gene repertoires highlights a complex and redundant molecular resistance architecture.

**Tables Supplementary 3–5** show strong associations between resistance phenotypes and dense repertoires of β-lactamase and efflux genes across species. In *Klebsiella pneumoniae*, MDR/XDR isolates were dominated by *bla_KPC* with frequent coexistence of *bla_NDM* and *CTX-M* variants. In *Acinetobacter baumannii*, carbapenem resistance was primarily linked to *OXA-type* carbapenemases. In *Pseudomonas aeruginosa*, resistance was characterized by widespread *mex* efflux operons with additional carbapenemase determinants, highlighting a redundant, polygenic resistance architecture in ICU isolates.

### Whole-Genome Resistome and Virulome Profiling

**Table 2** and **Table Supplementary 6** summarize whole genome resistome profiles across susceptible, MDR, and XDR isolates. Progressive expansion of resistance gene repertoires was observed with increasing resistance phenotypes. MDR/XDR *Klebsiella pneumoniae* showed enrichment of aminoglycoside resistance genes (p = 0.009), whereas *Acinetobacter baumannii* displayed significant increases in β-lactamases (p = 0.008), aminoglycoside (p = 0.007), and other resistance genes determinants (p = 0.011). In *Pseudomonas aeruginosa*, differences were observed for β-lactamase (p = 0.040) and fluoroquinolone-associated genes (p = 0.009).

**Table 2-.**
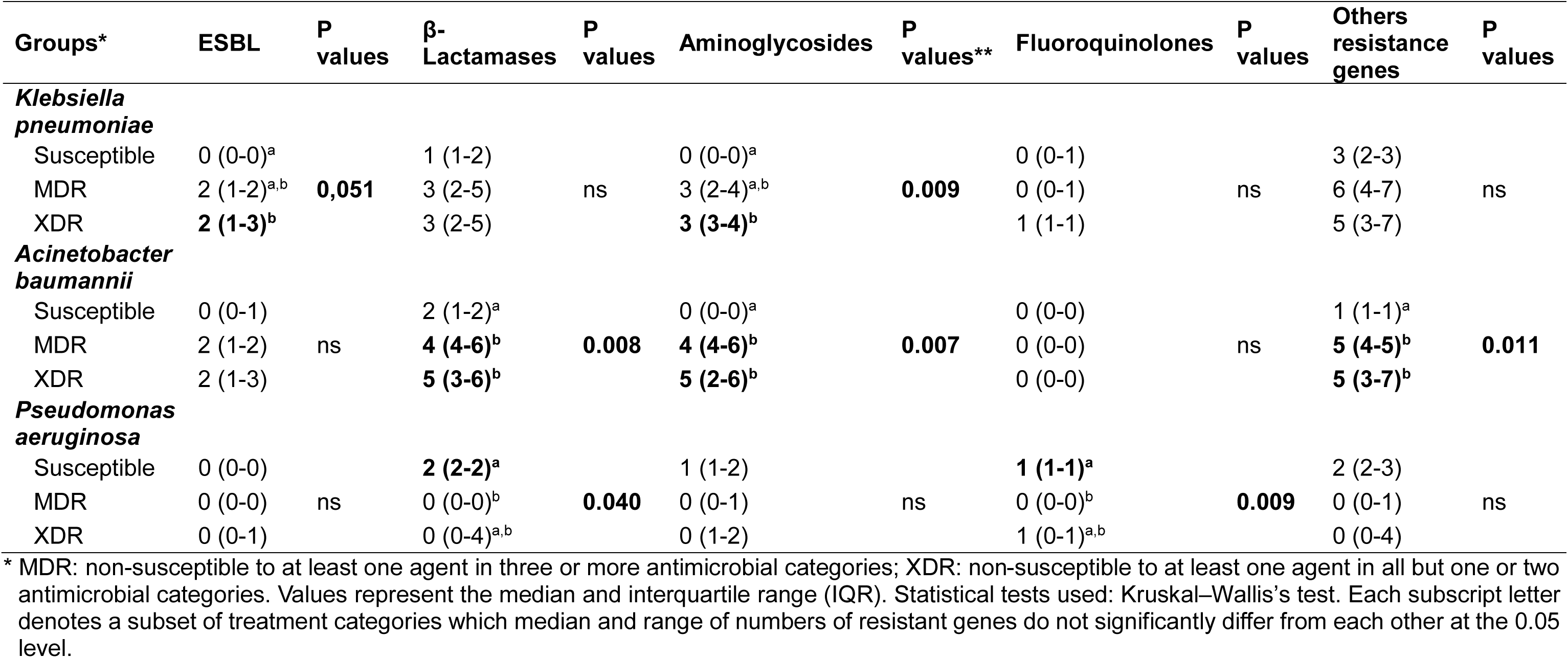
Comparative quantitative analysis of whole-genome resistance gene repertoires across susceptible, MDR, and XDR phenotypes in *Klebsiella pneumoniae*, *Acinetobacter baumannii*, and *Pseudomonas aeruginosa* isolates from an ICU in Northeast Brazil (2019–2021).

**Tables Supplementary 7** and **8** summarize whole-genome virulome profiles across susceptible, MDR, and XDR isolates available data in *K. pneumoniae* and *P. aeruginosa*. Virulence gene repertoires remained largely conserved across resistance phenotypes, with no consistent reduction in virulence determinants among MDR or XDR strains. These findings indicate preservation of pathogenic potential despite the accumulation of antimicrobial resistance mechanisms in ICU-associated Gram-negative pathogens.

### Proteomic Remodeling Across Resistance Phenotypes in ICU Gram-Negative Pathogens

Comparative label-free quantitative proteomic analyses identified phenotype-associated differences across susceptible (SUSC), multidrug-resistant (MDR), and extensively drug-resistant (XDR) isolates of *Klebsiella pneumoniae*, *Acinetobacter baumannii*, and *Pseudomonas aeruginosa* (**Figures 3–5**). Volcano plot analyses (|log₂FC| > 1; p < 0.05) showed differential abundance of proteins related to outer membrane components, transport systems, regulatory proteins, and metabolic processes across resistance phenotypes.

**Figure 3.**
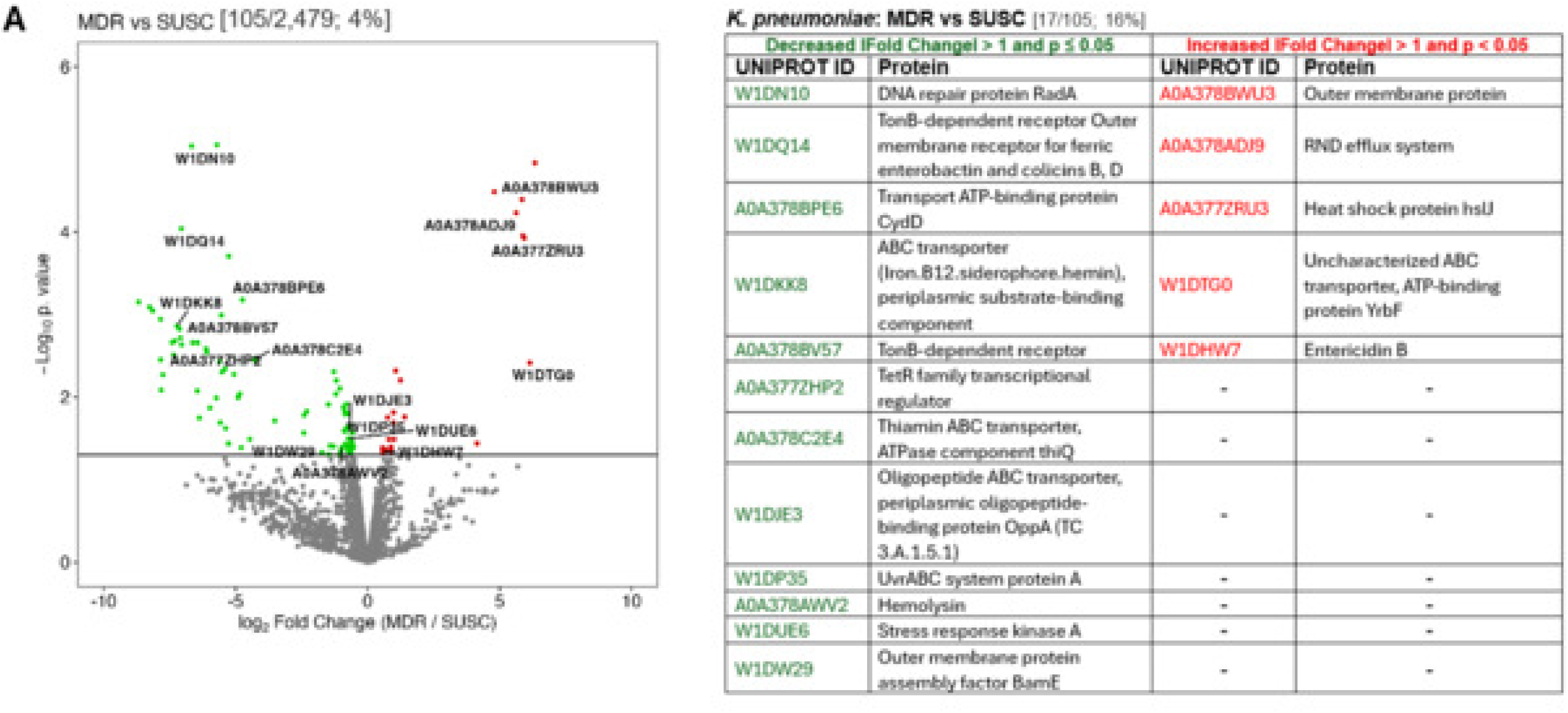
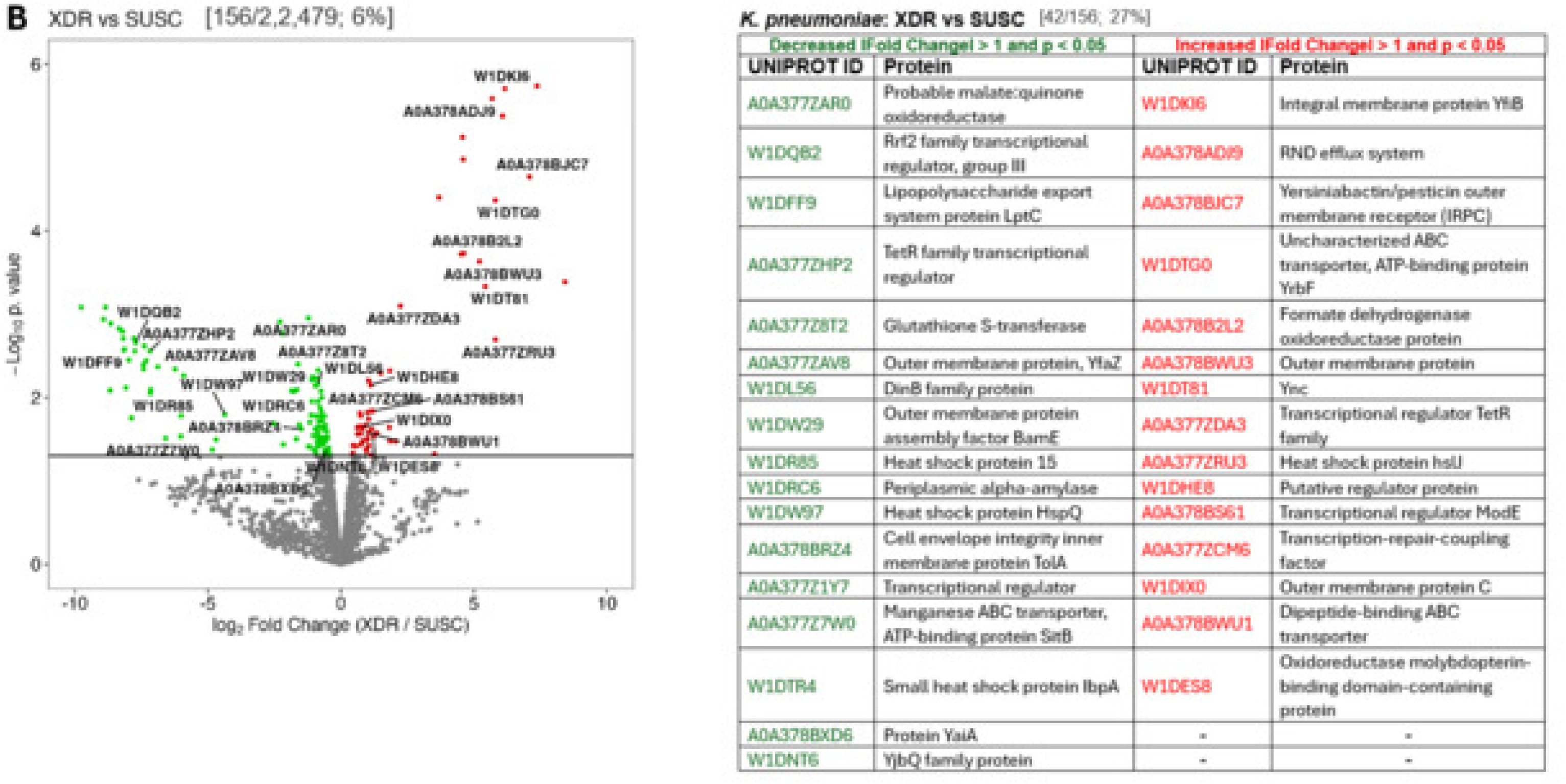
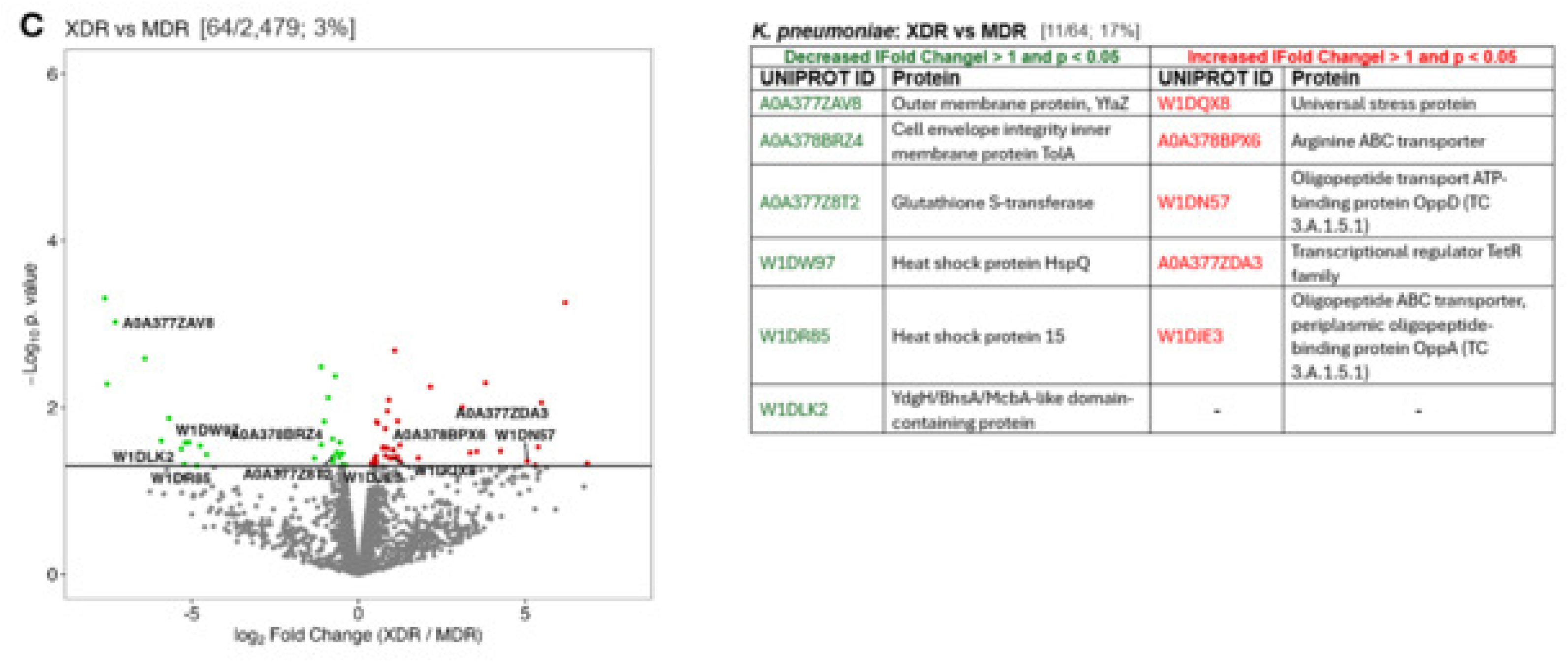
Differential protein abundance among resistance phenotypes of *Klebsiella pneumoniae*. Volcano plots showing pairwise comparisons of protein abundance among antimicrobial resistance phenotypes of *Klebsiella pneumoniae* isolates analyzed by label-free LC–MS/MS proteomics. Proteins are plotted according to log₂ fold change (x-axis) and −log₁₀ p-value (y-axis). Proteins achieving significance criteria (|fold change| > 1 and p < 0.05) are highlighted. (**A**) Multidrug-resistant (MDR) versus susceptible (SUSC) isolates; (**B**) Extensively drug-resistant (XDR) versus susceptible (SUSC) isolates; and (**C**) XDR versus MDR isolates. Each dot represents an individual quantified protein. Red dots indicate increased abundance, and green dots indicate decreased abundance relative to the comparison group.

**Figure 4.**
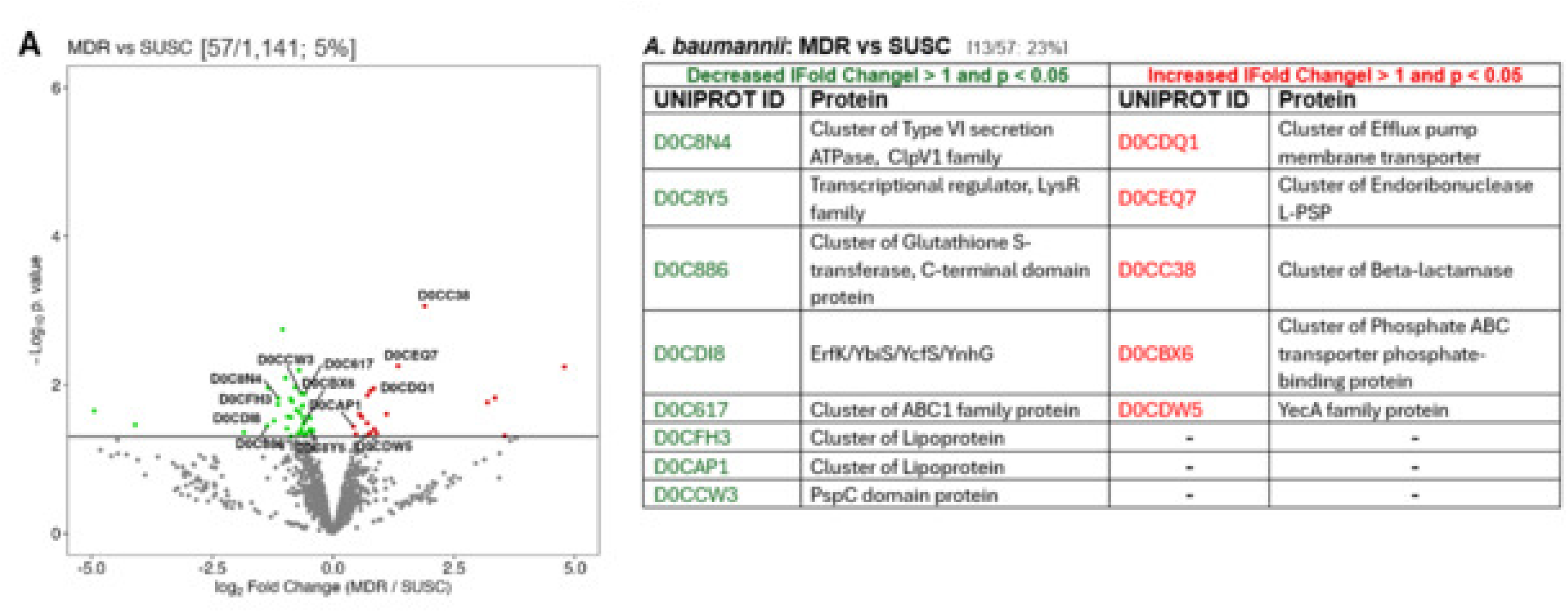
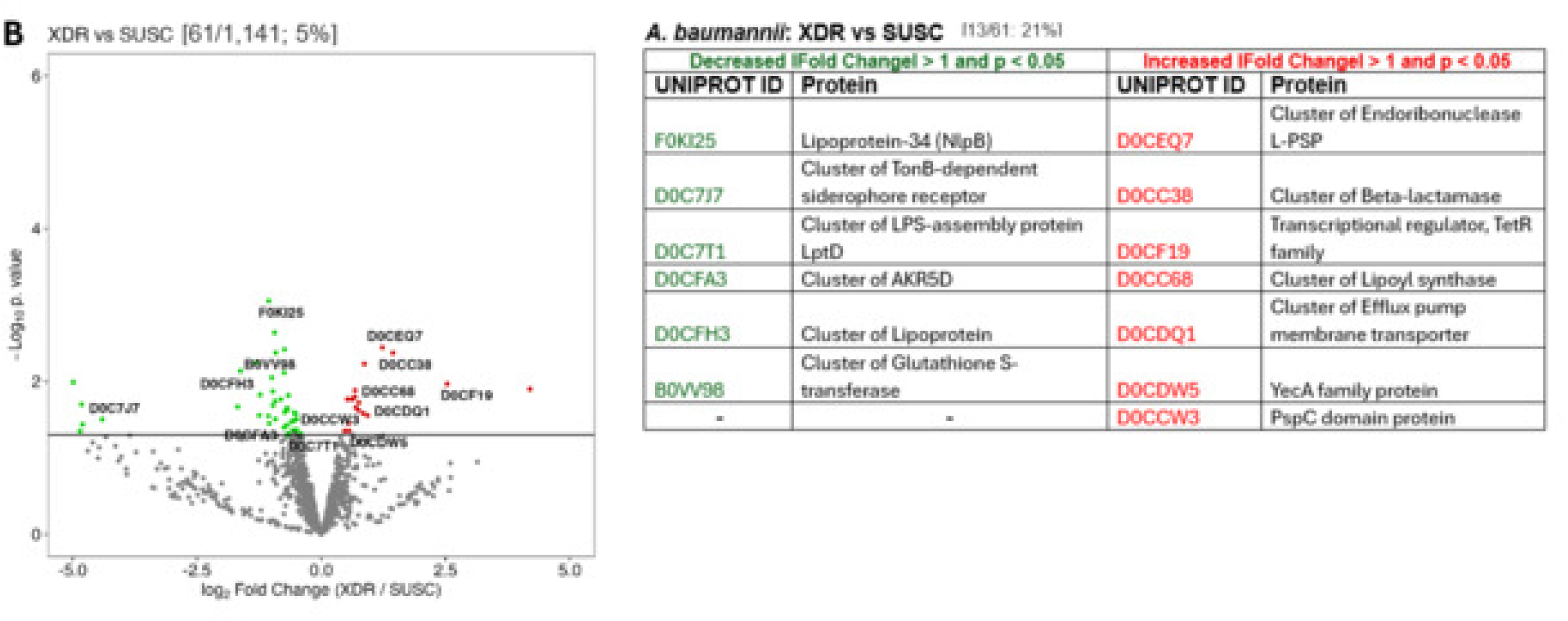
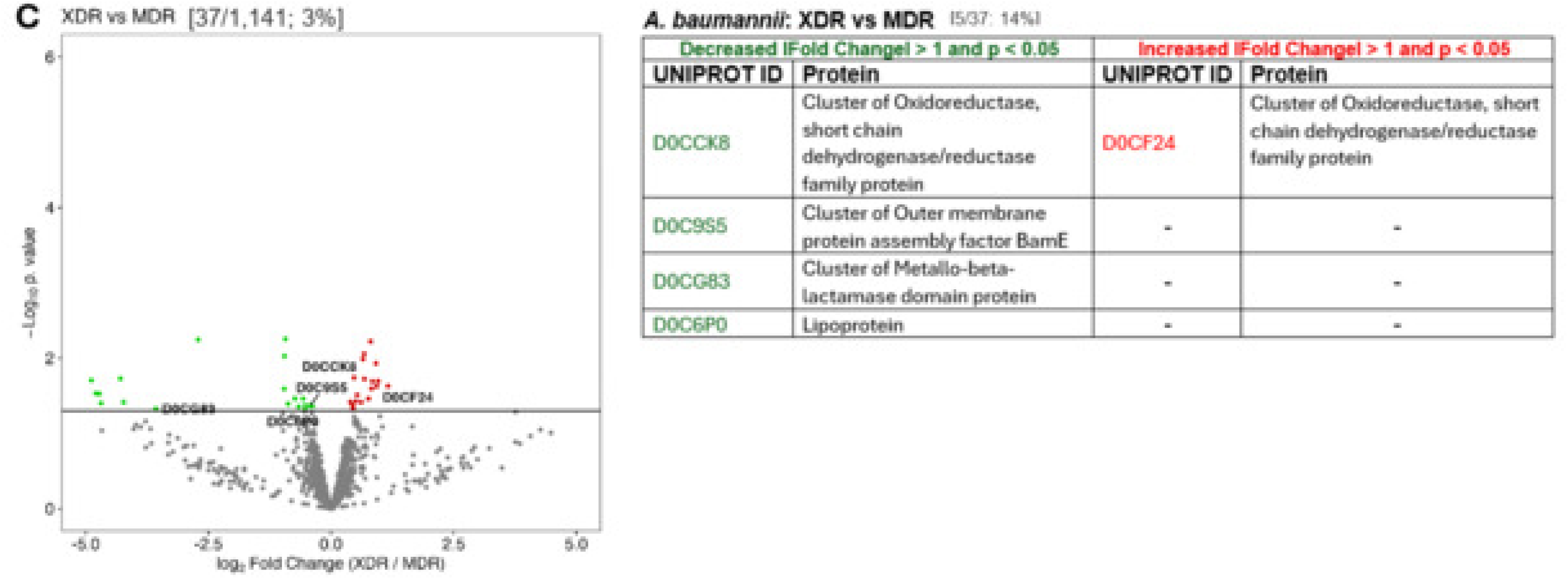
Volcano plot analysis of proteomic differences in *Acinetobacter baumannii* resistance phenotypes. Volcano plots displaying differential protein abundance among antimicrobial resistance phenotypes of *Acinetobacter baumannii* isolates determined by label-free quantitative LC–MS/MS analysis. Proteins are represented by log₂ fold change (x-axis) and −log₁₀ p-value (y-axis). Statistical significance was defined as |fold change| > 1 with p < 0.05. (**A**) MDR versus susceptible (SUSC) isolates; (**B**) XDR versus susceptible (SUSC) isolates; and (**C**) XDR versus MDR isolates. Each point corresponds to a detected protein. Proteins with increased abundance are shown in red and those with decreased abundance in green.

**Figure 5.**
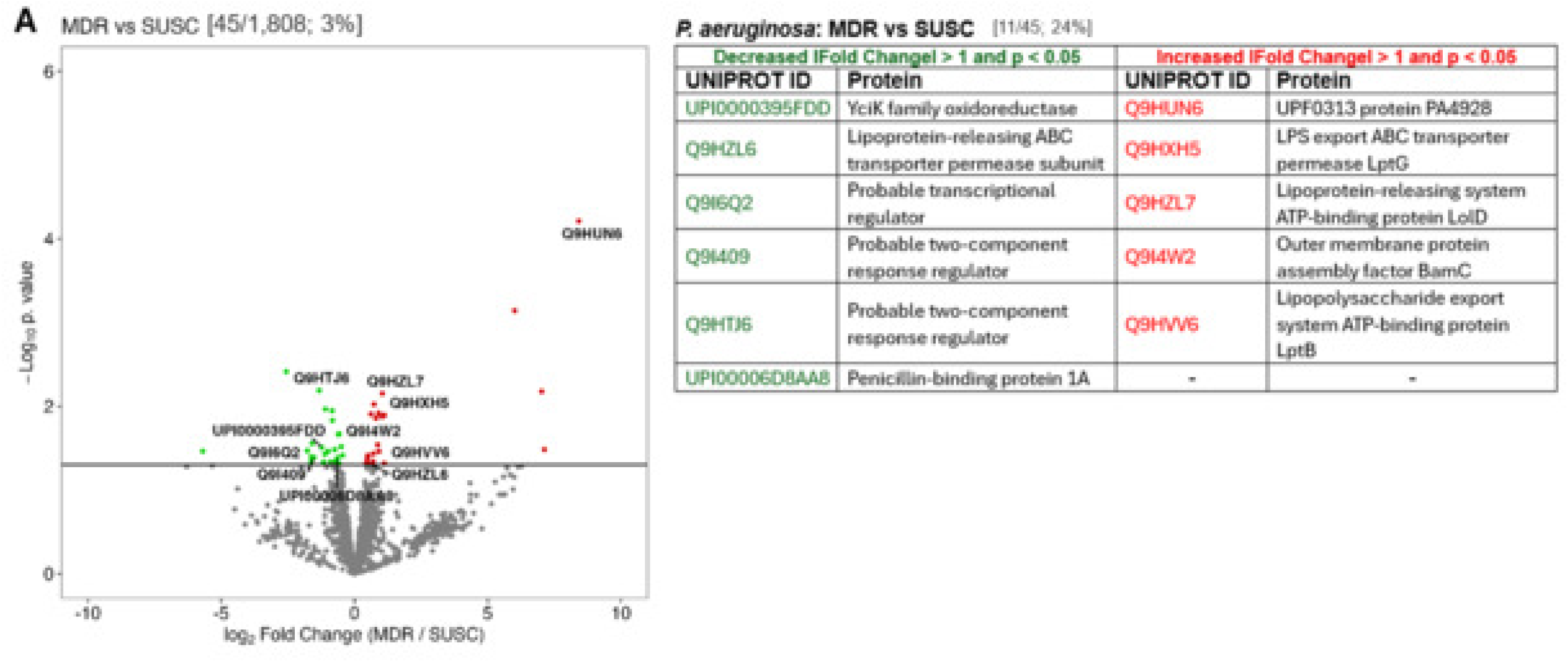
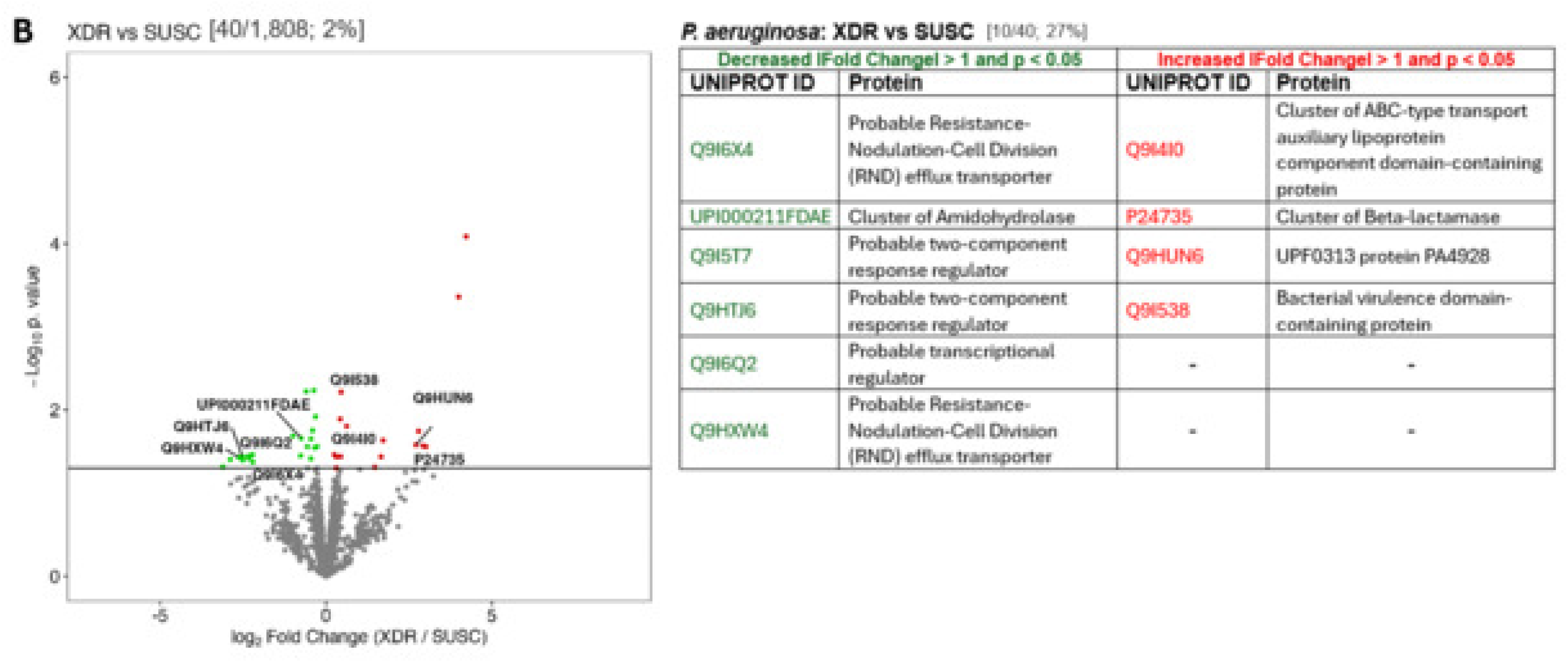
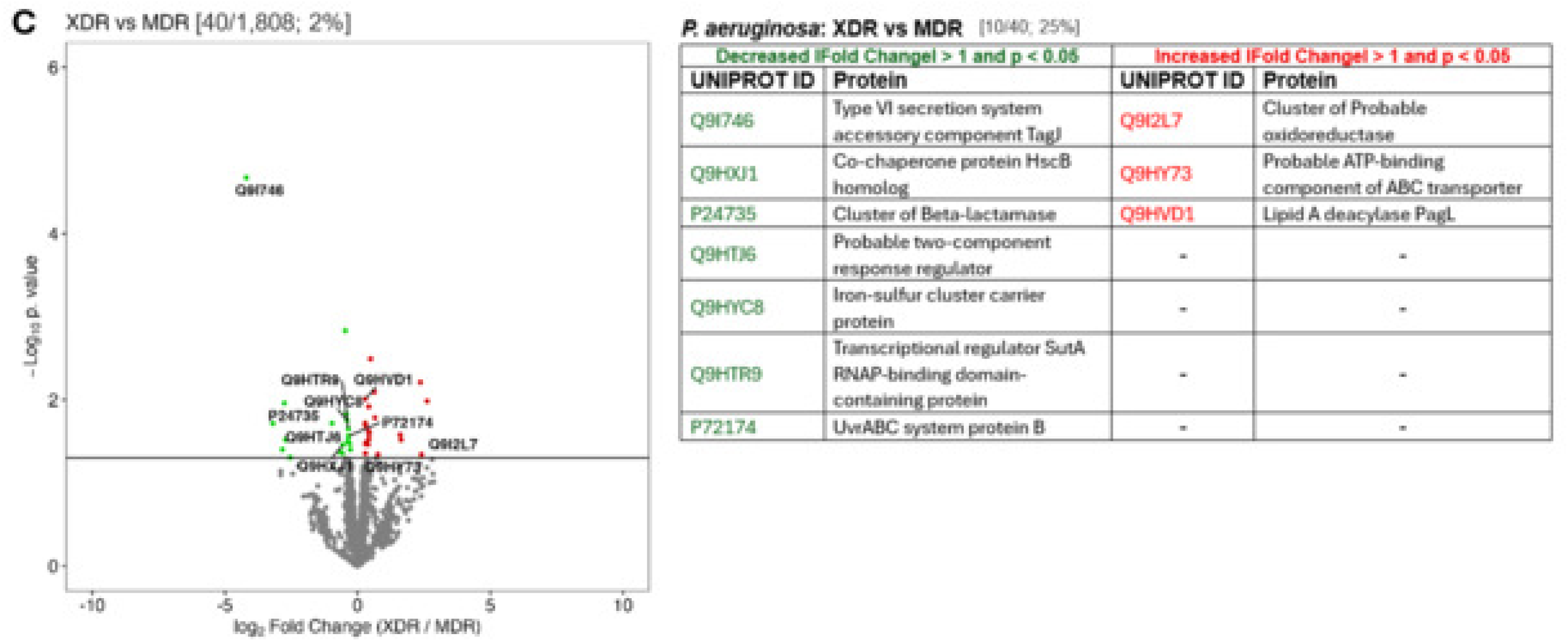
Proteomic comparisons across resistance phenotypes of *Pseudomonas aeruginosa*. Volcano plots illustrating pairwise comparisons of protein abundance among antimicrobial resistance phenotypes of *Pseudomonas aeruginosa* isolates analyzed using label-free LC–MS/MS proteomics. The x-axis represents log₂ fold change, and the y-axis represents −log₁₀ p-value. Proteins were considered significantly different when |fold change| > 1 and p < 0.05. (**A**) MDR versus susceptible (SUSC) isolates; (**B**) XDR versus susceptible (SUSC) isolates; and (**C**) XDR versus MDR isolates. Each dot represents one quantified protein; red indicates increased abundance and green indicates decreased abundance relative to the comparator phenotype.

In *K. pneumoniae*, MDR isolates showed increased abundance of outer membrane-associated proteins and transport-related systems, including RND efflux components and heat-shock protein HslJ (**Figure 3A**), while proteins involved in iron uptake (TonB-dependent receptors), ABC transport, and DNA repair (RadA) were reduced compared with SUSC isolates. XDR isolates demonstrated increased abundance of additional membrane-associated proteins, including YfiB, YrbF, and TetR-family regulators (**Figure 3B**), alongside reduced levels of proteins such as LptC, BamE, glutathione S-transferase, and small heat-shock proteins. In comparisons between XDR and MDR isolates (**Figure 3C**), differences were observed in proteins including universal stress proteins, OppD, and TetR-family regulators.

In *A. baumannii*, MDR isolates exhibited increased abundance of membrane transporters, β-lactamase-associated proteins, and phosphate ABC transporter components relative to SUSC strains (**Figure 4A**), whereas proteins such as LysR-family regulators, glutathione S-transferases, and lipoproteins showed lower abundance. XDR isolates displayed increased abundance of β-lactamase-associated proteins, TetR-family regulators, and transport-related proteins (**Figure 4B**), with reduced levels of TonB-dependent receptors, LptD, and lipoproteins. Comparisons between XDR and MDR isolates (**Figure 4C**) showed differences in oxidoreductase-related proteins, BamE, and metallo-β-lactamase domain proteins.

In *P. aeruginosa*, MDR isolates showed increased abundance of proteins associated with lipopolysaccharide transport (LptB, LptG), lipoprotein transport (LolD), and outer membrane assembly (BamC) (**Figure 5A**), with reduced levels of penicillin-binding protein 1A and selected regulatory proteins. XDR isolates demonstrated increased abundance of β-lactamase-associated proteins and ABC transporter components (**Figure 5B**), while some efflux-associated and regulatory proteins showed lower abundance. In XDR versus MDR comparisons (**Figure 5C**), differences were observed in ABC transporter ATP-binding proteins, PagL, TagJ, iron–sulfur cluster proteins, and SutA.

Overall, differences in protein abundance across SUSC, MDR, and XDR phenotypes were observed in proteins related to membrane structure, transport systems, regulatory proteins, and metabolic functions (**Figure 6**).

**Figure 6.**
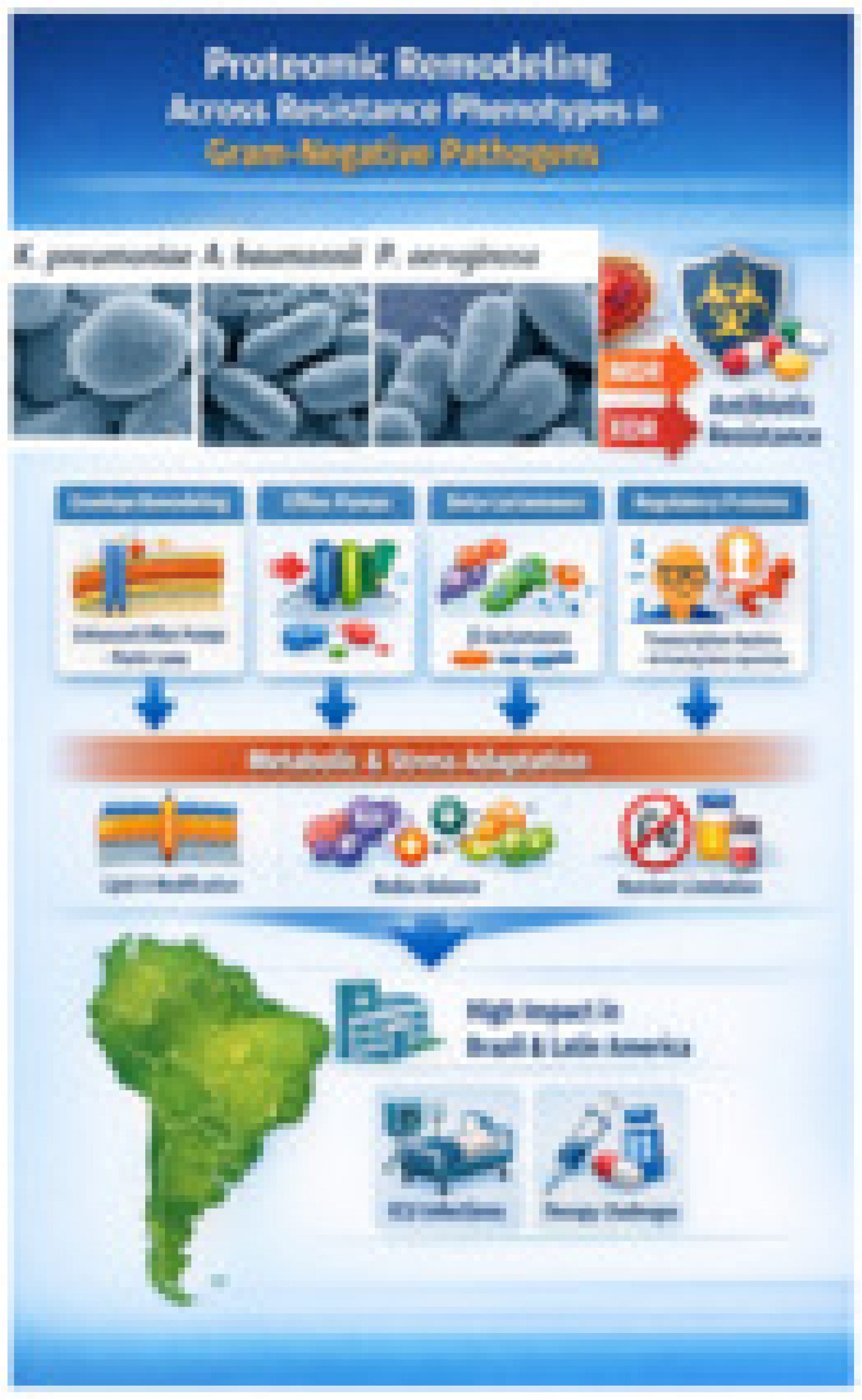
Graphical summary of proteomic remodeling across resistance phenotypes in Gram-negative pathogens. Conceptual graphical summary illustrating the study design and main proteomic features identified across antimicrobial resistance phenotypes of *Klebsiella pneumoniae*, *Acinetobacter baumannii*, and *Pseudomonas aeruginosa*. The schematic depicts the comparison of multidrug-resistant (MDR) and extensively drug-resistant (XDR) isolates and summarizes protein functional categories represented in the proteomic dataset, including envelope-associated proteins, efflux pump systems, β-lactamases, regulatory proteins, and proteins related to metabolic and stress-associated processes. The lower section presents the clinical context addressed in the study, highlighting intensive care unit infections and therapeutic challenges in Brazil and Latin America. This figure provides a visual overview of the analytical framework and thematic organization of the proteomic findings.

## Discussion

This study provides an integrated phenotypic, genomic, and proteomic characterization of Gram-negative pathogens circulating in an ICU in Northeastern Brazil. *Klebsiella pneumoniae*, *Acinetobacter baumannii*, and *Pseudomonas aeruginosa* were the predominant species and exhibited a high burden of antimicrobial resistance, including elevated rates of MDR and XDR phenotypes. These findings are consistent with global reports identifying these organisms as priority pathogens associated with adverse outcomes in critically ill patients (3–5,15,16).

High resistance rates to β-lactams, carbapenems, aminoglycosides, and fluoroquinolones were observed, consistent with ICU environments where antimicrobial pressure, invasive procedures, and prolonged hospitalization contribute to resistance selection (7,15–16). Carbapenem-resistant *A. baumannii* has been associated with high mortality in pulmonary and bloodstream infections (4,5,17), and the high prevalence of carbapenemase-producing *K. pneumoniae*, particularly KPC- and NDM-producers, aligns with reports from Latin America and globally (13,18–20). These findings are consistent with antimicrobial resistance surveillance data in Brazil (6).

Genomic analyses revealed widespread coexistence of β-lactamase genes across species. In *K. pneumoniae*, bla_KPC was highly prevalent, frequently co-occurring with bla_NDM and bla_CTX-M variants, consistent with genomic surveillance studies (12,13,18). In *A. baumannii*, OXA-type carbapenemases, particularly bla_OXA-23 and bla_OXA-51, predominated (4,5,18,21). In *P. aeruginosa*, resistance involved both β-lactamases and intrinsic mechanisms, including efflux systems (8,9,22,23). The near-universal detection of RND efflux operons and the presence of acrAB-tolC in *K. pneumoniae* and *P. aeruginosa* support the contribution of efflux-mediated resistance across species (9,22,24,25), with species-specific differences observed in *A. baumannii* (21,22,24).

Whole-genome analyses showed expansion of resistance gene repertoires from susceptible to MDR and XDR phenotypes without consistent reduction in virulence gene content. This preservation of virulence determinants alongside increasing resistance has been reported in previous genomic studies (18,20,23,26) and is relevant for ICU settings.

Proteomic analyses identified phenotype-associated differences in proteins related to membrane structure, transport systems, regulatory proteins, and metabolic processes. Across species, MDR and XDR isolates showed differential abundance of outer membrane-associated proteins, transporters, and regulatory elements compared with susceptible isolates. These findings are consistent with prior genomic and transcriptomic studies of Gram-negative pathogens (11,13,18,19,27,28).

In *K. pneumoniae* and *A. baumannii*, MDR and XDR phenotypes were associated with differences in proteins linked to membrane structure, transport systems, and β-lactamase-related functions, alongside reduced abundance of proteins involved in nutrient acquisition. In *P. aeruginosa*, differences included proteins associated with lipopolysaccharide transport, membrane assembly, and β-lactamase-related components. Increased representation of transcriptional regulators, including TetR-family proteins, was also observed across species. These patterns are consistent with reported mechanisms involving efflux systems, porin changes, and β-lactamase activity (3,5,9,10,22,29,30).

Taken together, these findings indicate that antimicrobial resistance in ICU-associated Gram-negative pathogens involves the coexistence of multiple resistance genes and phenotype-associated proteomic differences. The integration of genomic and proteomic data complements conventional susceptibility testing and provides additional resolution for characterizing resistance phenotypes.

From a clinical perspective, the high prevalence of MDR and XDR phenotypes and the coexistence of multiple resistance mechanisms highlight the importance of rapid diagnostics and optimized antimicrobial strategies in ICU settings. The presence of β-lactamases and efflux-associated systems supports current recommendations for β-lactam/β-lactamase inhibitor combinations and other targeted therapies (3–5,9,17). At the same time, variability in resistance-associated mechanisms underscores the need for continued surveillance and individualized treatment approaches.

This study has limitations, including its single-center design, which may limit generalizability. However, the integration of phenotypic, genomic, and proteomic data provides a comprehensive characterization of high-priority pathogens in a critical-care setting. In conclusion, Gram-negative pathogens isolated from ICU patients exhibited high levels of antimicrobial resistance characterized by the coexistence of multiple resistance genes and phenotype-associated proteomic differences. The progression from susceptible to MDR and XDR phenotypes was associated with expansion of the resistome without loss of virulence determinants. These findings highlight the complexity of antimicrobial resistance in ICU settings and support the use of integrative approaches for improved characterization and surveillance.

## Author Contributions

A.A.M.L., D.T.S., and A.H. conceptualized and designed the study. M.S.M., L.F.B.N., M.A.F.C., S.A.R., F.P.M., L.F.B.L., A.E.S.A., A.C.D., L.M.V.C.M., and R.N.D.G. conducted bibliographic review, sample processing, and coordinated data collection. M.S.M., L.F.B.N., M.A.F.C., A.H., I.F.N.L., J.L.N.R., L.V.C.F., D.T.S., N.E.S., F.M., B.M.C., É.A.G.A., and A.A.M.L. developed the study methodology and experimental framework. M.S.M., L.F.B.N., D.T.S., N.E.S., and M.A.F.C. performed genomic and bioinformatic analyses. N.E.S., D.T.S., M.S.M., M.A.F.C., F.M., and A.A.M.L. performed and interpreted proteomic analyses. M.S.M., L.F.B.N., J.K.S., S.A.R., F.P.M., and L.F.B.L. conducted in vitro validation experiments and laboratory testing. J.Q.S.F., D.V.S.C., F.S.J., and A.A.M.L. performed statistical analyses and data interpretation. M.S.M., L.F.B.N., and A.A.M.L. drafted the manuscript. All authors (including D.T.S., N.E.S., A.H., I.F.N.L., J.L.N.R., L.V.C.F., D.V.S.C., S.A.R., F.P.M., J.K.S., L.F.B.L., A.E.S.A., A.C.D., L.M.V.C.M., R.N.D.G., F.M., B.M.C., and É.A.G.A.) contributed to critical revision of the manuscript for important intellectual content and approved the final version. A.H., D.T.S., D.V.S.C., and A.A.M.L. supervised the study. A.A.M.L. administered the project and acquired funding.

## ETHICAL APPROVAL

Ethics approval by the Brazilian National Research Ethics Committee, (n° 03300218.2.0000.5054), statement (n° 3.143.258).

## DATA AVAILABILITY STATEMENT

All data supporting the findings of this study are included in the manuscript. Whole-genome sequencing and proteomic datasets are available from the corresponding author upon reasonable request.

## ACKNOWLEDGEMENT

The article processing charge (APC) for this publication was funded by the Coordenação de Aperfeiçoamento de Pessoal de Nível Superior (CAPES) (ROR identifier: 00×0ma614). To open access, the authors have applied a Creative Commons Attribution (CC BY) license to any Author Accepted Manuscript arising from this submission. During the preparation of this manuscript, the authors used ChatGPT Plus (version 5.2, OpenAI) and Paperpal Prime (Editage, Cactus Communication Service Pte Ltd, Singapura) as an editorial assistance tool to support language refinement, structural organization, clarity improvement, and formatting of sections of the text. The AI tool was not used to generate original scientific data, perform analyses, interpret results, or create figures or tables. All experimental design, data collection, statistical analyses, data interpretation, and scientific conclusions were developed and verified exclusively by the authors. The authors critically reviewed and edited all AI-assisted content and took full responsibility for the accuracy, integrity, and originality of the manuscript.

